# Single-character insertion-deletion model preserves long indels in ancestral sequence reconstruction

**DOI:** 10.1101/2024.03.09.584071

**Authors:** Gholamhossein Jowkar, Jūlija Pěcerska, Manuel Gil, Maria Anisimova

## Abstract

Insertions and deletions (indels) play a significant role in genome evolution across species. Realistic modelling of indel evolution is challenging and is still an open research question. Several attempts have been made to explicitly model multi-character (long) indels, such as TKF92, by relaxing the site independence assumption and introducing fragments. However, these methods are computationally expensive

On the other hand, the Poisson Indel Process (PIP) assumes site independence but allows one to infer single-character indels on the phylogenetic tree, distinguishing insertions from deletions. PIP’s marginal likelihood computation has linear time complexity, enabling ancestral sequence reconstruction (ASR) with indels in linear time. Recently, we developed ARPIP, an ASR method using PIP, capable of inferring indel events with explicit evolutionary interpretations

Here, we investigate the effect of the single-character indel assumption on reconstructed ancestral sequences on mammalian protein orthologs and on simulated data. We show that ARPIP’s ancestral estimates preserve the gap length distribution observed in the input alignment. In mammalian proteins the lengths of inserted segments appear to be substantially longer compared to deleted segments. Further, we confirm the well-established deletion bias observed in real data

To date, ARPIP is the only ancestral reconstruction method that explicitly models insertion and deletion events over time. Given a good quality input alignment, it can capture ancestral long indel events on the phylogeny

## 1 Introduction

Insertion and deletion (indel) events produce significant amounts of natural variation in species genomes. Consequently, indels make a major contribution to complex evolutionary processes. Today indel variants in genomic sequences can be reliably documented and studied due to improvements in sequencing methods. In closely related species, differences attributed to indels (per base pair) are several-fold more frequent than substitution events (Britten et al, 2003; Wetterbom et al, 2006). In the human genome, up to a quarter of all genomic variants are due to indels, most of which are very short (Mills et al, 2006). While indels are distributed across both coding and non-coding parts of genomes, they are far more frequent in non-coding sequences. Compared to substitutions, indel changes are expected to have a stronger deleterious effect on functional proteins (Tóth-Petŕoczy and Tawfik, 2013), also explaining their lower prevalence in coding sequences. Despite this, many deleterious coding indel variants persist in the human population and can cause disease-related gene defects (e.g., (Chuzhanova et al, 2003)).

In comparative studies of sequence evolution, indels are represented as gaps in alignments of homologous sequences. With growing divergence, different indel events can merge and overlap, masking the mutational history. Nevertheless, alignment gaps carry much phylogenetic information (Dessimoz and Gil, 2010), which can provide valuable insights for evolutionary studies when analyzed correctly. However, properly modelling the evolutionary process of insertions and deletions is challenging from the computational and modelling perspective, and there is no gold standard in the field. In fact, many evolutionary studies either completely ignore indels, or heavily trim indel-rich sequence regions, due to the lack of software tools implementing appropriate models. Disentangling individual insertion and deletion events based on the observed gap distributions in a multiple sequence alignment (MSA) requires stochastic models of sequence evolution that also include the insertion and deletion processes over time. Substitutions are traditionally described via Markov models, which assume site independence, while indels violate this assumption since each indel event can involve multiple residues. Therefore, models properly including these events tend to be computationally expensive.

The first evolutionary model with indels, TKF91, lifted the assumption of site independence and described single-character indels via a birth-death process (Thorne et al, 1991). As TKF91 models single-character events, it implies a linear gap cost in the MSA inference, but due to the non-independence of sites, the complexity of computing the marginal likelihood under this model is exponential in the number of taxa. Bouchard-Côté and Jordan proposed the PIP model, a close relative of TKF91, where insertions follow the Poisson process while deletions are added to the Markov substitution model as an absorbing state. The complexity of marginal likelihood computation under the PIP model is reduced to linear, which allows for this model to be adopted for phylogenetic inferences (Zhai and Bouchard-ôté, 2017; Maiolo et al, 2018, 2021; Jowkar et al, 2023). However, like TKF91, PIP explicitly models only single-character indels.

Modelling longer indels as several independent single-character events lacks biological realism and could lead to biases such as homology histories with too many events, alignments with scattered gaps, and high indel rates. Some evolutionary indel models allow long indels (Thorne et al, 1992; Mikĺos et al, 2004; De Maio, 2021). For example, the TKF92 model, an extension of TKF91, is also a birth-death process but with indels happening as unbreakable multiple-site fragments with a geometric length distribution (Thorne et al, 1992). This modelling assumption, however, means that TKF92 cannot explain overlapping indels. The “long indel” model (Mikĺos et al, 2004) relaxed the unbreakable fragment assumption but assumed infinite sequences. Both these models can be considered an approximation of the Generalised Geometric Indel (GGI) model (Holmes, 2020). However, while the lengths of individual indels have a geometrical distribution, the length distribution of observed gaps in the alignment is not geometric in general. Considering that models with long indels also tend to be computationally slow, these are currently of little practical value for large datasets.

Computationally, PIP holds promise for practical phylogenetic analyses despite the single-character indel assumption. For example, we showed that PIP-based alignment inference can pick up multiple-character indels (long indels) when the data strongly suggests this (Maiolo et al, 2018, 2021). Zhai and Bouchard-ôté demonstrated that modelling indel evolution and indel rate variation improves the accuracy of phylogeny reconstruction when using the PIP model and its generalizations.

Recently, we proposed a PIP-based ancestral sequence reconstruction (ASR) approach implemented in ARPIP (Jowkar et al, 2023). Apart from Bayesian MCMC implementations (e.g., Historian (Holmes, 2017)), ARPIP is the only ASR method that uses an explicit model of indel evolution and can infer the specific locations of insertions and deletions on the tree. Another popular ASR method is FastML-webserver (Ashkenazy et al, 2012), which uses the so-called “indel-coding” method to include indels. This approach does not include a proper statistical model of insertion and deletion and implies that a deleted character can be reinserted. GRASP (Ross et al, 2022), another recent method, accommodates indels in the ASR inference by representing sequences as partial order graphs. However, as with indel-coding, deleted characters can be reinserted, and there is no explicit model governing the indel process.

## 2 The goals of this study

Having an explicit model of indel evolution is desirable; however, an over-simplistic model could also have a detrimental effect on the resulting inferences, including over-estimation of indel rates and scattered ancestral sequence alignments by including too many single-character gaps. Therefore, we aim to investigate whether using the single-character indel assumption negatively impacts ASR. Since ASR methods typically take a fixed MSA and phylogeny as input, using good-quality input MSAs and phylogenetic trees is imperative for accurate ASR, irrespective of the method used. While MSA quality is still quite an elusive concept in general, here we assume that a good-quality MSA captures multiple-character (long) indels in a phylogenetically consistent way. Therefore, in our study, we use PRANK (Löytynoja and Goldman, 2005), the phylogeny-aware tool which infers phylogenetically meaningful gaps by distinguishing insertions from deletions in a progressive manner on the tree.

Here, given accurate input data, we assess the systematic bias in PIP-based ASR by investigating the fragmenting of gaps in the inferred sequences at the ancestral nodes of the phylogeny. To test this, we present a large-scale analysis of protein orthologs from six mammalian species (human, three primates, and two rodents), taken from the popular orthologous protein database OMA (Altenhoff et al, 2021), as well as analysis of simulated data. We chose this specific phylogenetic dataset for two reasons. First, the mammalian species tree for these specific taxa is unambiguous and can be accepted as “true” (although the indel history is unknown, see (Nichols, 2001)). Second, insertion and deletion biases in these species have long been a subject of interest, meaning that our findings can be interpreted in the context of current literature. For these data, we evaluated per-site insertion and deletion frequencies in different lineages and compared the gap distributions in the observed and inferred sequences.

To get a better understanding of ASR properties and potential biases under PIP, we proceed by analyzing simulated data. In our simulations, we mimic the OMA-based protein orthologous groups so that the results on real data can be compared to expected performance on very similar data where the truth is known. Our results suggest no significant difference in observed and inferred ancestral gap length distributions. This means that ARPIP tends to preserve the long indels from the input alignment in the inferred ancestral sequences. We also could confirm the well-documented deletion bias (Zhang and Gerstein, 2003; Ogurtsov et al, 2004; Tao et al, 2007; Lin et al, 2017; He et al, 2019; Loewenthal et al, 2021).

## 3 Results

### 3.1 Results on mammalian data

We extracted and analyzed 12*^′^*088 orthologous protein groups, each containing one sequence from six *eutherian* mammals. Sequences in each orthologous group were aligned, and ancestral sequences were reconstructed given the inferred multiple sequence alignment (MSA) and the species tree (see data and methods; Fig. 12). For each site in an MSA, our ASR method ARPIP infers the most likely insertion and deletion history, allowing us to distinguish insertion and deletion events. Note that the reconstruction is done independently for each site, as in all other ASR methods. Therefore, we evaluated the number of inserted and deleted residues per site and per time interval rather than counting multiple residue events. This way of measuring indel rates is intuitively similar to substitution rates; therefore, it has a simple interpretation without having to account for the length of the full indel. Another advantage of this approach is that it makes it easy to evaluate the impact of indel events on sequence length over time. Note that here, we make a clear distinction between gaps and indels. Namely, gaps are the stretches of missing characters (gap characters “-”) in a sequence resulting from the alignment. In contrast, indels are insertion and deletion events inferred via the ancestral reconstruction with ARPIP. Gaps in MSAs can appear due to several multiple-character insertions and deletions. Since ASR is performed independently at each site, gaps spanning multiple sites are described as a series of single-character indel events at several affected individual sites. To evaluate whether this assumption is reasonable during ASR, we study whether the ARPIP method preserves the distribution of gap lengths of the input MSA in the sequences reconstructed at ancestral nodes.

#### 3.1.1 Comparing the number of inserted and deleted characters

1*^′^*267 of orthologous groups had no gaps in the inferred MSAs, presumably due to strong conservation. These groups were therefore excluded from the indel statistics presented here. For the remaining 10*^′^*769 orthologous groups, the total numbers of inserted and deleted residues on the species tree are visualized in Fig. 1, and more detailed statistics are presented in Tab. 1. The *human* lineage had the lowest number of inserted and deleted characters, as well as overall gap characters in the sequences (4.5% of total sequence length). This is strongly contrasted by the *gorilla* lineage, which experienced the highest indel numbers among all studied species with 6.2% of its total sequences in MSAs consisting of gaps. Only the *chimp* and the *macaque* lineages had more gaps in their sequences, with 6.5% and 6.8%, respectively. These three primate lineages also had the longest on average gap lengths (on average 14.8 amino acids for *chimp*, 15.2 for *gorilla* and 15.3 for *macaque*), compared to *human* (10.6) and all other lineages. *Hominini* ancestral lineage experienced the lowest number of inserted residues and indels overall, although this can be expected since this divergence corresponds to the shortest branch length on the species tree.

**Fig. 1.**
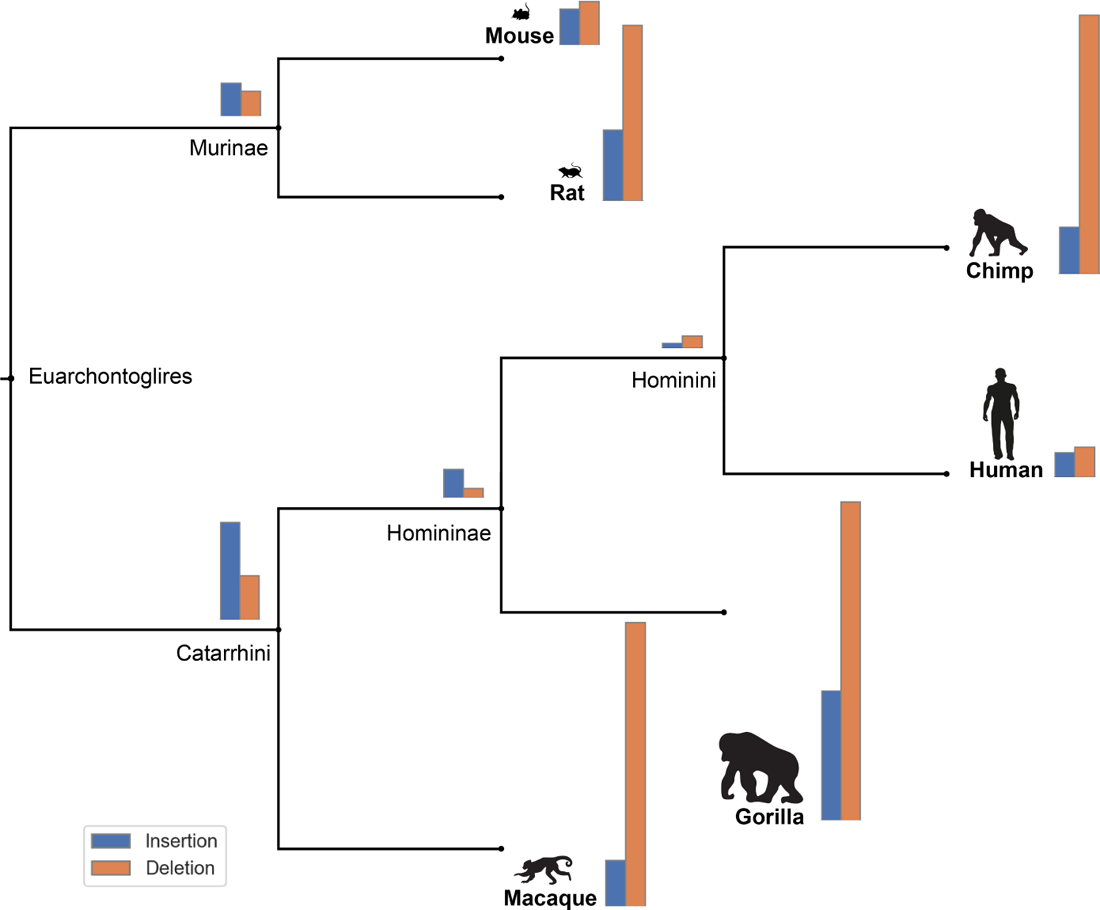
Number of indel events across studied species. *Gorilla* has the largest number of indel events per lineage while *Hominini* and *Homininae* have the lowest number of indel events, respectively (see Tab. 1).

**Table 1.**
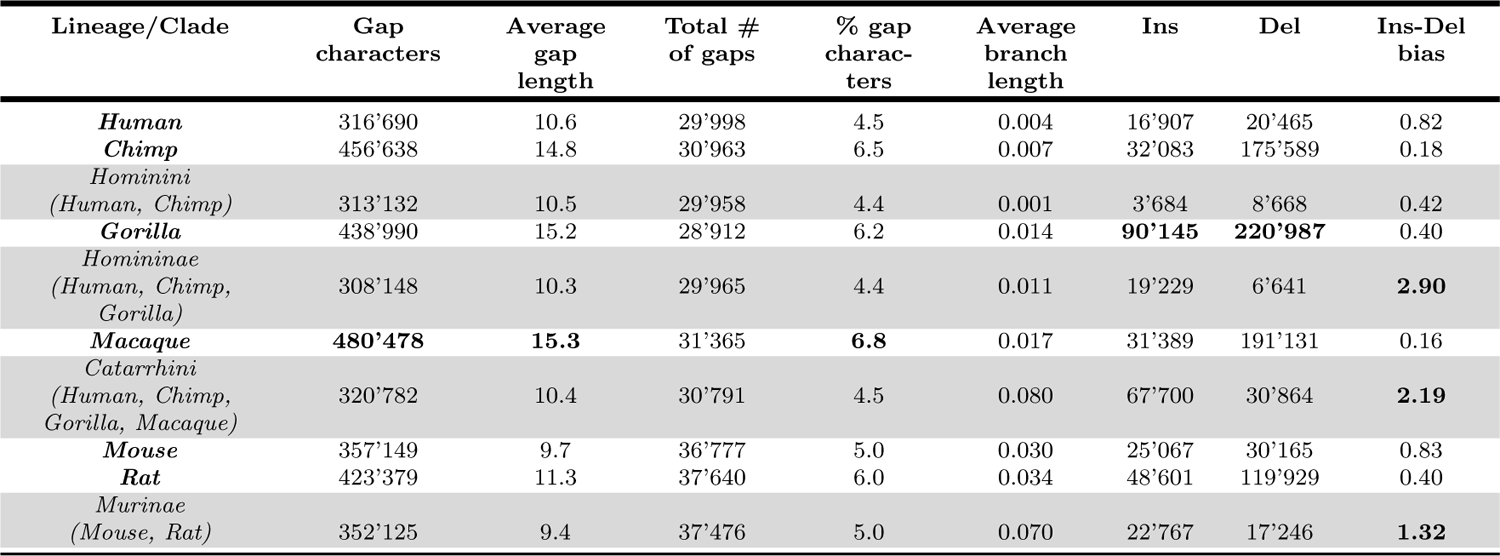
Summary statistics of gaps and indels on mammalian data.

Next, we calculated the insertion-deletion bias as the ratio between the numbers of insertion and deletion events (see Fig. A1 in Appendix). Overall, the number of deletions was larger than the number of insertions for all six extant lineages. The bias towards deletions was particularly strong in *macaque* (0.16) and *chimp* (0.18), but also well pronounced in *gorilla* and *rat* (both 0.41). In contrast, most ancestral lineages displayed a bias towards insertions, which was particularly pronounced in the *Homininae* (2.90) and *Catarrhini* ancestors (2.19).

#### 3.1.2 Tracing sequence lengths along the tree

Further, we investigated whether the observed deletion bias in extant lineages affects the sequence length dynamics across the species phylogeny. For each orthologous group, we computed Spearman correlation coefficients between sequence lengths (observed at the leaves or inferred at the ancestral nodes, gap characters removed) and the evolutionary distance from the root to its corresponding node in the tree (evolutionary age). The majority of analyzed orthologous groups showed no significant correlations at a 5% significance threshold. Nevertheless, we observed significant correlations in 12.04% of orthologous groups with positive correlations for 459 genes and negative correlations for 838 genes (Fig. 2). This suggests that 7.78% of analyzed gene sequences had the tendency to shrink, while 4.26% had shown a tendency to grow.

**Fig. 2.**
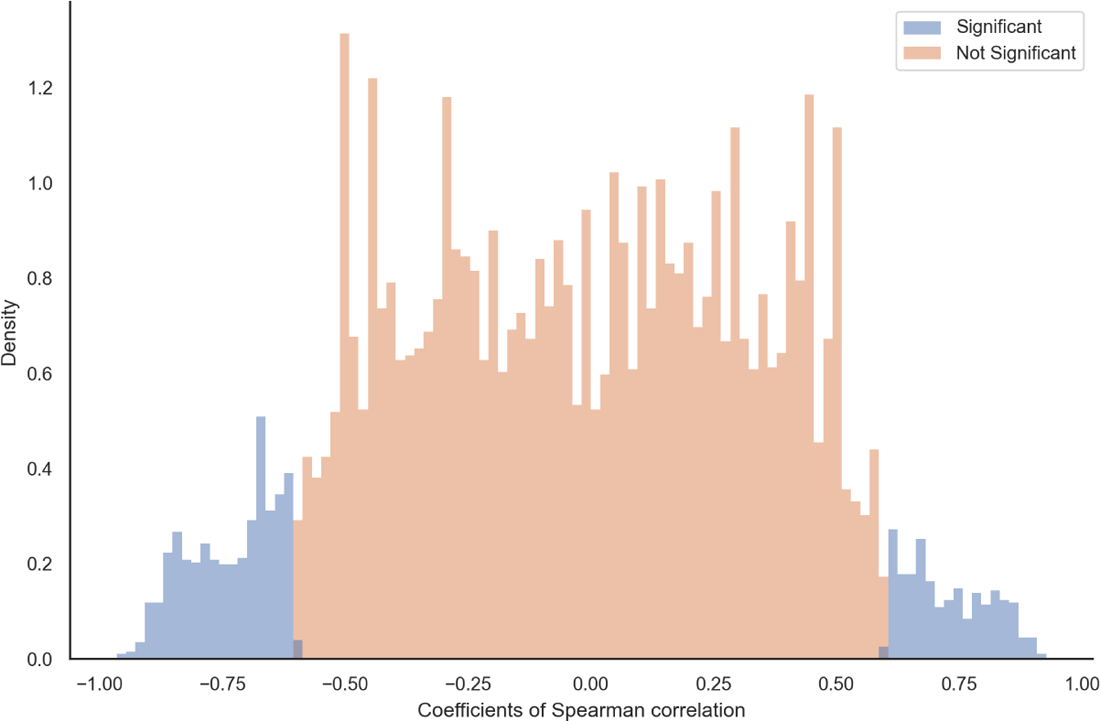
The distribution of Spearman correlation coefficients between sequence length (at the tips and root) and evolutionary distance from the root per OMA groups on six mammalian species.

#### 3.1.3 Gap length distribution preserved over time

We asked whether the gap distributions in the six observed sequences differed from those in the inferred ancestral sequences. The gap distribution in the inferred MSA of the six observed sequences results from the PRANK alignment and would, therefore, exhibit any inherent systematic biases of the PRANK method, if any. By analyzing whether a change in gap length distribution occurs at the inferred ancestral sequences, we aim to evaluate whether ARPIP tends to bias the distribution in a given alignment towards shorter gaps.

Such an effect is expected to be maximal at the root of the tree. Therefore, we compared the empirical distribution of gap lengths at the root with the distribution at the leaves over all OMA groups. Visually, the two distributions match (Fig. 3).

**Fig. 3.**
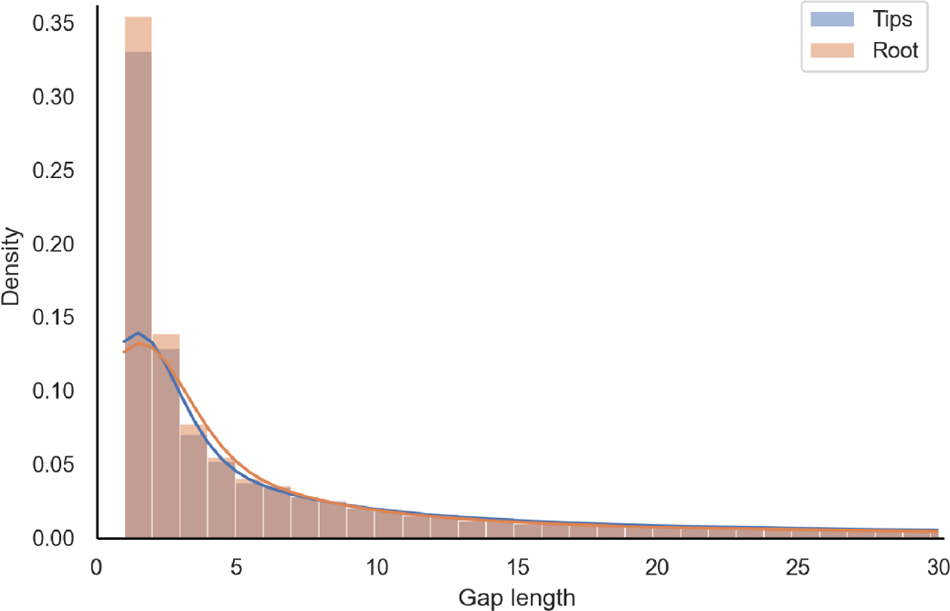
The empirical gap length distribution of tips vs. root on mammalian sequences. The plot is the histogram with 100 bins, while the upper bound of gaps is limited to 100 residues.

Furthermore, for each OMA group, we computed the mean gap lengths at the root and the mean gap lengths at the tips. The differences between the means are distributed around zero with a heavier tail in the positive range, which leads to an average difference of 4 characters, meaning that gaps at the tips tend to be around 4 characters longer (Fig. 4).

**Fig. 4.**
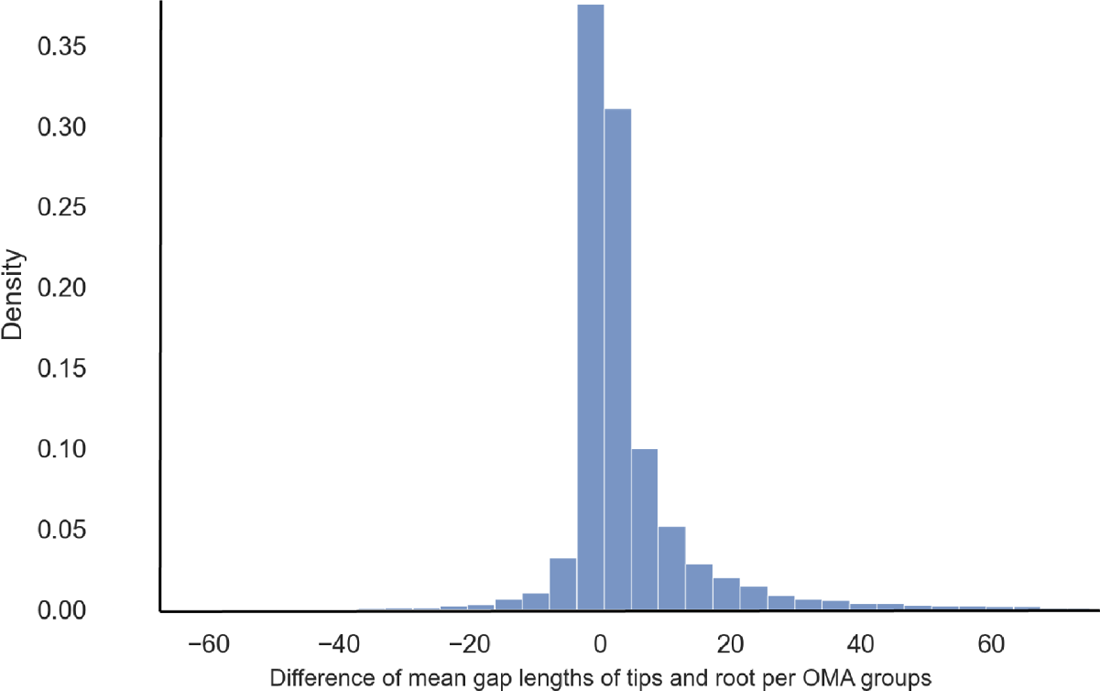
Paired difference of mean gap lengths per OMA groups on mammalian data (with 100 bins).

#### 3.1.4 Inserted segments are longer than deleted segments

Finally, we compared the empirical distributions of multiple-character insertion and deletion events over time on the phylogeny. Fig. 5 depicts that the empirical distributions of insertions and deletions are consistent with the empirical gap length distribution as single-character events are the most frequent, and their frequency decreases as the length of the event increases. In addition, we observed that insertions tend to be significantly longer than deletions, where the mean insertion length was 17.81, while it was 8.67 for deletion events.

**Fig. 5.**
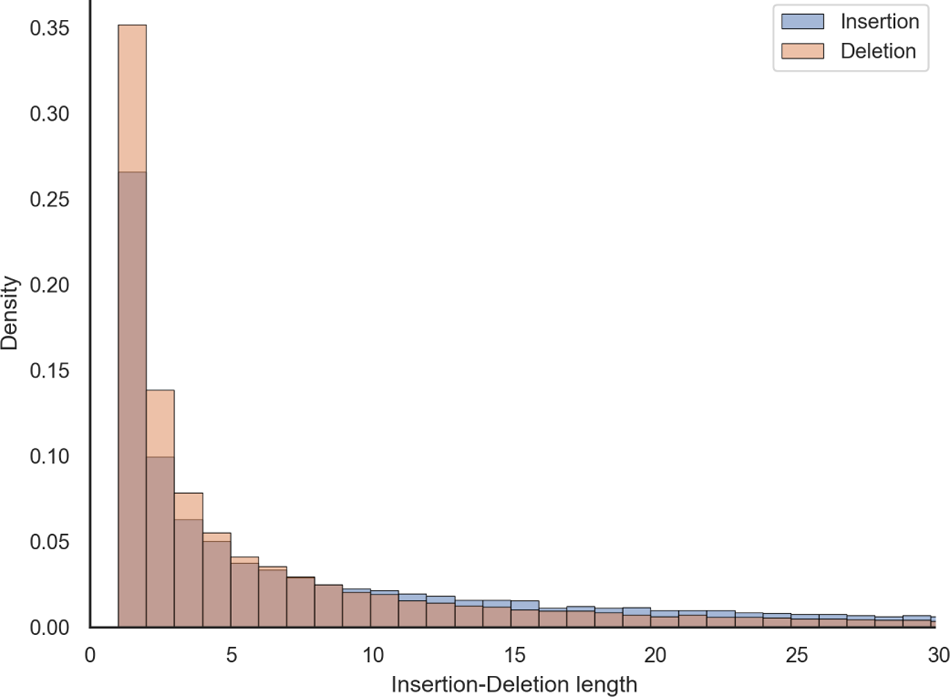
The empirical distribution of inserted vs. deleted segment lengths. The plot is the histogram with 100 bins, while the upper bound of gaps is limited to 100 residues.

### 3.2 Results on simulated data

To study ARPIP under fully controlled conditions, we have simulated sequences with INDELible. To set realistic parameters, we sampled 1000 random OMA groups. For each sampled OMA group, we used the corresponding PhyML tree to evolve a replicate on it, with the root sequence length of 1000 amino acids, indel rate of 0.1, and indel lengths distributed according to the Zipfian distribution with exponent 1.7. INDELible’s maximum indel length parameter was set to the length of the longest gap in the PRANK MSA of the OMA group in question. We supplied the true simulated MSA of the observed sequences to ARPIP for all the analyses.

#### 3.2.1 Reconstruction accuracy

On simulated data, ARPIP inferred a positive insertion-deletion bias in all nodes of the trees; i.e., more individual characters were inserted than deleted (Appendix Fig. A2). It correctly reconstructed more than 98% of ancestral residues, resulting in 90% correctly inferred ancestral columns (Appendix Tab. A1). The average precision^1^ in gap character inference was 94%, with a recall^2^ of 97%. We divided the simulation results according to the F-score (a measure of predictive performance defined as the harmonic mean of precision and recall) in gap retrieval into “optimal” (132 samples with F-score *≥* 99%) and “sub-optimal” (858 samples with F-score *<* 99%). Fig. 7 shows the branch length distributions for the two classes. Note that for the sub-optimal samples, this distribution is near zero. For these samples, we observed a lower accuracy in gap reconstruction. Indeed, shorter branches provide less information, and we expect larger variances and lower accuracy. Likewise, this happens also for extremely high divergences when the signal becomes saturated. Furthermore, the insertion probability in PIP is proportional to branch lengths. Thus, the choice of insertion points also depends on the relative branch lengths of the phylogeny. Fig. 6 shows the ROC curve points for each sample (and not just one point, i.e. the average).

**Fig. 6.**
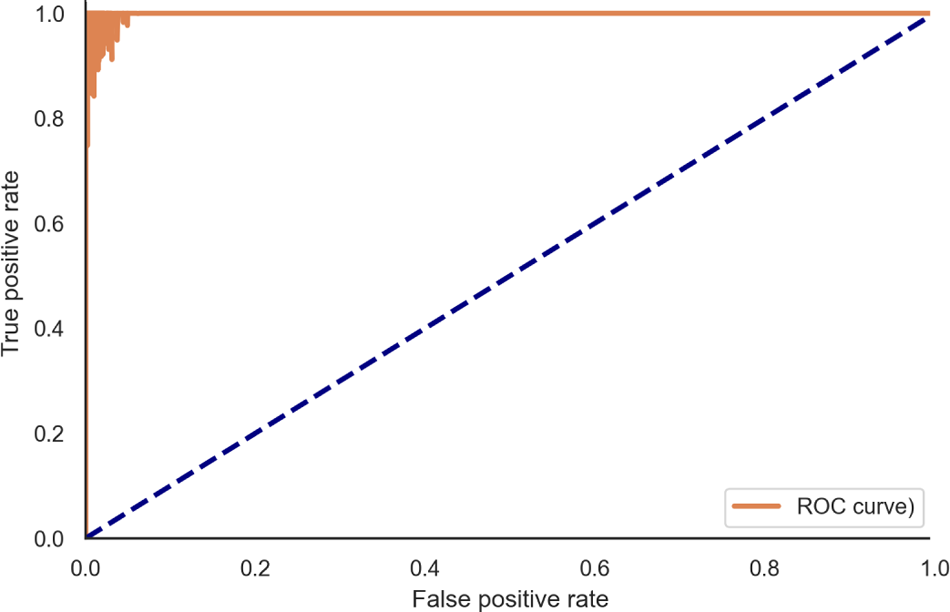
ROC curve: true positive (recall or sensitivity) vs. false positive (1-specificity) rates at the ARPIP gap estimation.

**Fig. 7.**
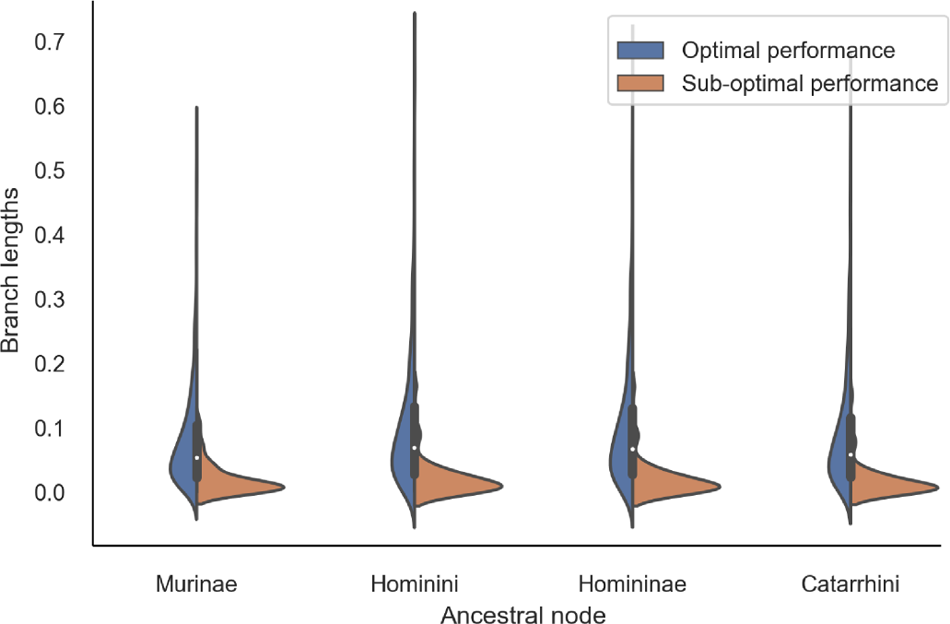
Distribution of ancestral node branch lengths in the simulated data, grouped by inference performance.

#### 3.2.2 Tracing sequence lengths along the tree

Analogous to the real data analysis above, we correlated the sequence length without gaps in each node with the node’s evolutionary age for each replicate. Again, the majority of the Spearman coefficients were not significant at the threshold of 5%. Among the 14.95% significant ones, we observed 93 positive and 55 negative correlations (Fig. 8). Contrary to the real data, here, the majority of the significant replicates tended to grow, while 37% were shrinking. This is consistent with the positive indel bias.

**Fig. 8.**
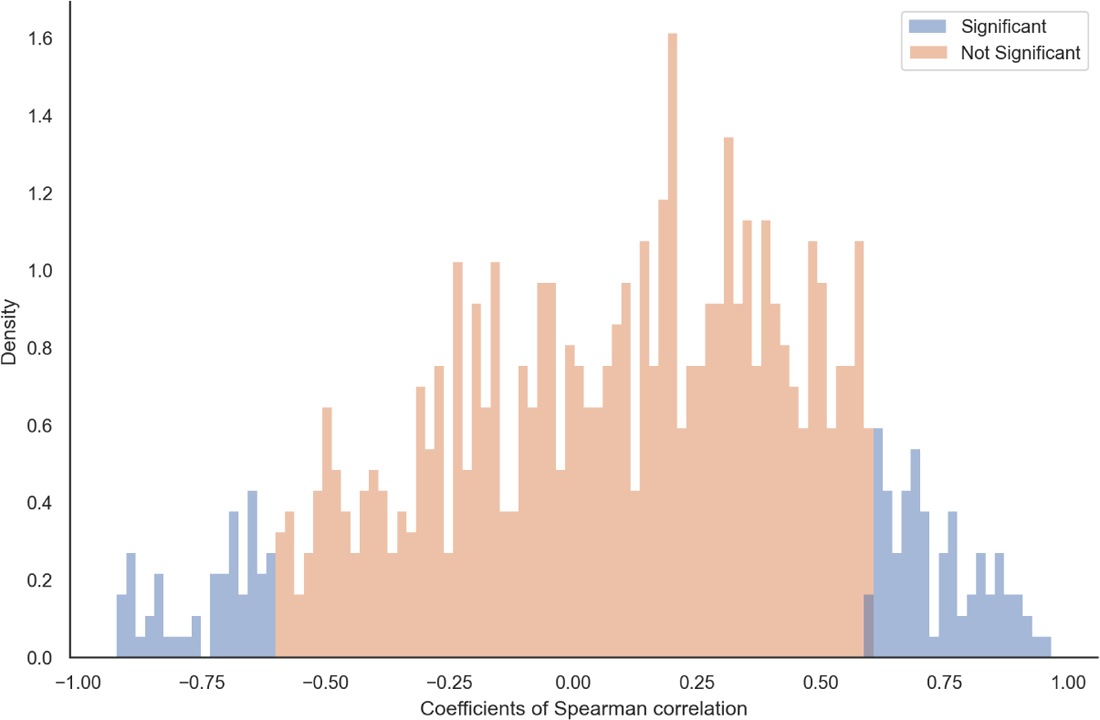
The distribution of Spearman correlation coefficients between sequence length (at the tips and root) and evolutionary age per OMA group on simulated data.

#### 3.2.3 Gap length distribution is preserved over time

Next, we asked if the gap length distribution in the inferred ancestral sequences differed from the true distribution, i.e. the one generated by the simulation. The two distributions match (Fig. 9) and have a Kullback-Leibler divergence of 4.42 *×* 10*^−^*^5^. According to the PIP model, we expect sequence lengths to be preserved, meaning not shrinking nor growing. Furthermore, there seems to be no decline of gap lengths towards the root of the tree, as the gap length distribution inferred at the root of the tree matches the distribution in the observed sequences at the leaves (Fig. 10). Note that in contrast to the real data case above, where the gaps at the leaves were inferred by PRANK, here we were able to compare to the true (simulated) MSA.

**Fig. 9.**
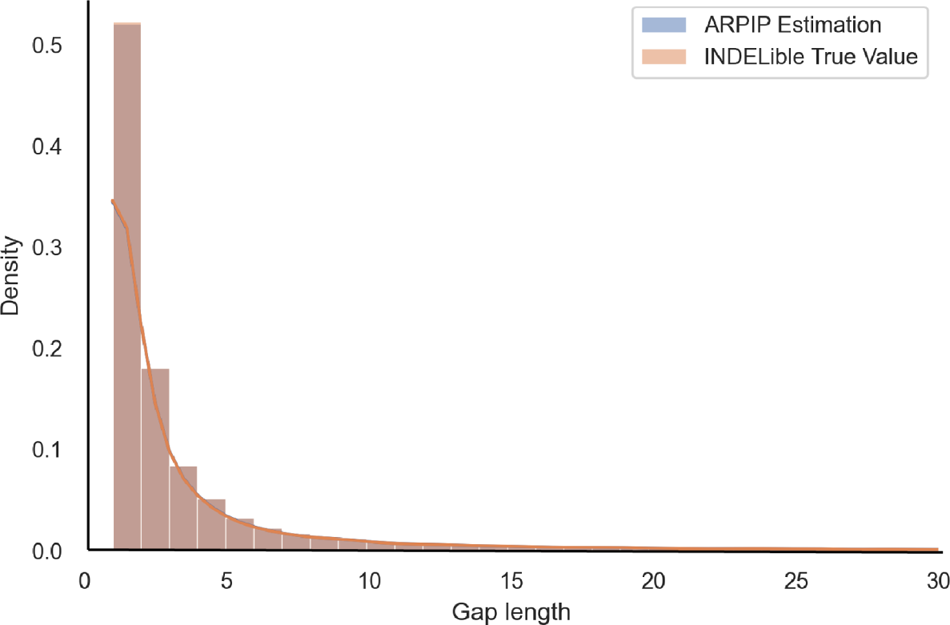
Overlapped distributions of gap lengths from ARPIP inference and INDELible true values.

**Fig. 10.**
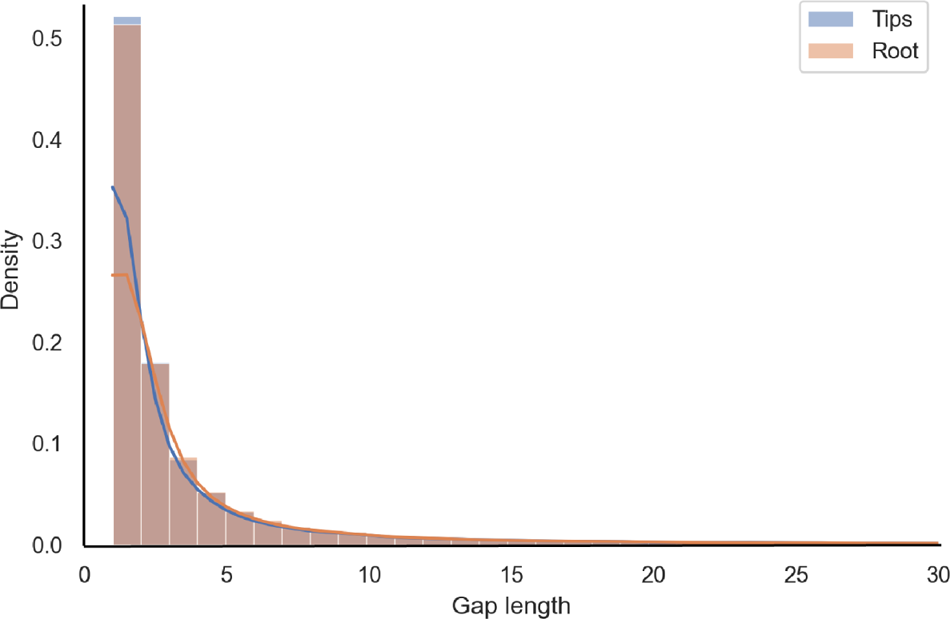
Empirical gap length distribution at the tips vs. the root in simulated sequences as a histogram with 100 bins with a cut-off at 100.

To further quantify the difference between simulated and inferred distributions, we computed the mean gap lengths at the root and the mean gap lengths at the tips for each of the 1000 replicates. Analogously to the mammalian data, the differences between the means were symmetrically distributed around zero (Fig. 11). The differences were not statistically different from zero (Mann-Whitney test, *p* = 0.67; two-sample t-test, *p* = 0.997). In summary, our simulation findings corroborate the results from real data. ARPIP preserves the gap lengths from the input alignment.

**Fig. 11.**
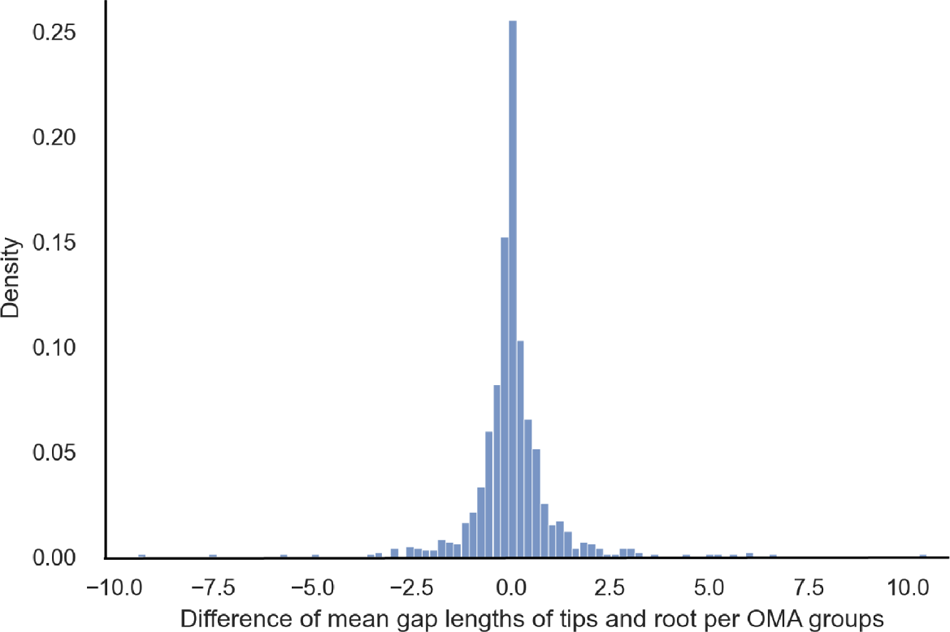
Paired difference of mean gap lengths per OMA groups on mammalian data (with 100 bins).

**Fig. 12.**
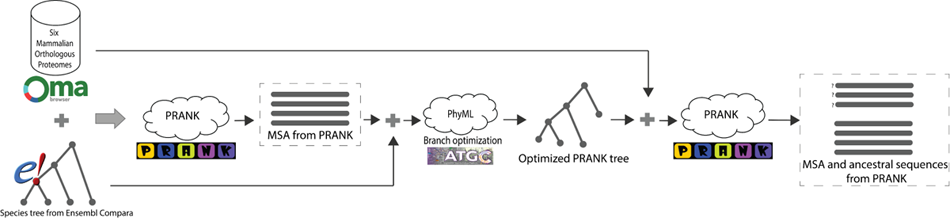
Algorithmic data acquisition pipeline.

## 4 Discussion and conclusions

Until recently, state-of-the-art ASR methods focused on inferring ancestral characters. Indels were often mishandled – either by removing gappy MSA columns, treating gaps as ambiguous characters (Yang, 2007), or reconstructing ancestral gaps with ad-hoc indel methods like “indel coding” (Ashkenazy et al, 2012). Further, such methods typically do not easily distinguish between insertions and deletions. Unlike previous approaches, ARPIP reconstructs insertions and deletions independently and uses the evolutionary indel model PIP. However, PIP only describes single-character indels.

In contrast to ASR, methods for MSA inference are more advanced with respect to allowing for long indels. One of the most advanced aligners is PRANK; it uses the phylogeny to distinguish insertions from deletions and, thus, infers phylogenetically meaningful long indels. All current ASR methods take an MSA as input. Here, we have shown on real data (with PRANK alignments) and by simulation (with the true simulated MSAs from INDELible) that the ancestral estimates by ARPIP preserve the long indel structure present in the MSA. This surprising result can partly be explained by the fact that under PIP the insertion and deletion points of a site only depend on the gap patterns (i.e. the presence and absence of gaps), and are independent of the character states (Jowkar et al, 2023). Neighboring sites with identical gap patterns form long indels and lead to identical indel histories (see, for example, Appendix B). Further studies will be needed to quantify how differences in neighboring gap patterns affect long indel preservation. Based on ARPIP’s strong performance, we hypothesize that minor pattern differences will still preserve most long indels. Also, analogously to other phylogeny-related inference problems (Bergsten, 2005) our simulations showed that short branches lead to a lower accuracy in gap reconstruction.

Furthermore, in line with the biology (de Jong and Rydén, 1981) and previous bioinformatics studies (Kuo and Ochman, 2009; Zhang and Gerstein, 2003; Loewenthal et al, 2021), we found that deletions are more frequent than insertions. Such deletion bias has been detected across the whole tree of life and has multiple possible evolutionary explanations. For example, (He et al, 2019) suggests that even strictly balanced insertion and deletion rates result in a linearly increasing genome size through time rather than a completely fixed genome size. The authors attribute this effect to the fundamental asymmetry of indels, as insertions produce more characters available for deletion, while deletions reduce the total number of characters, resulting in fewer deletable ones. The authors suggest that while the huge variety in genome sizes among species seems to require exponential size growth, the effective insertion bias cannot act for prolonged periods of evolutionary time. Consequently, the mechanisms producing larger genome sizes only act sporadically and are likely to be removed in the long term, making them very difficult to detect by looking into existing genomes. On the other hand, the commonly detected deletion bias could be an effect similar to the pull-of-the-present effect in phylodynamics, where younger lineages show seemingly higher birth/lower death rates (Nee et al, 1994). This effect stems from the fact that we are observing a snapshot of the evolutionary history that is cut off from the future, meaning that while some of the present-day lineages might go extinct, they have had less time to do so than older lineages and thus are more likely to have gotten sampled. In essence, this would mean that deletions might appear more frequently in the present sequences, but only because they have not yet been fixed in the underlying population at the moment of sampling.

Finally, until now, studies on indel length distributions have lumped the insertions and deletions together, often just inferring gap length distributions. As a step forward, we suggest inferring separate distributions for insertion and deletion lengths. Our findings from mammalian data strongly point to longer insertion lengths than deletion lengths. Further, given the higher prevalence of deletions and the remarkable uniformity of protein length distribution across the tree of life (Nevers et al, 2023), it is conceivable that the two distributions differ, with deletions lengths having a smaller mode than insertions. Recent work from Tal Pupko’s lab is a notable step in the direction of inferring indel length distributions based on event reconstruction (Wygoda et al, 2024).

## 5 Data and methods

### 5.1 Sequence acquisition and alignment

First, we used the OMA database (Altenhoff et al, 2021) to obtain orthologous protein sequences so that each orthologous group (OMA group) contained one sequence from each of six mammalian species, namely *human*, *chimp*, *gorilla*, *macaque*, *mouse*, and *rat*. The OMA database is known for its higher precision but lower recall compared with the majority of other methods (Altenhoff et al, 2019, 2021). A corresponding species tree was extracted from the Ensembl Compara v. 105 (Yates et al, 2020) by pruning a larger mammalian tree to the six species considered in this study (see Fig. 13). This species tree was then provided as a guide tree for reconstructing multiple sequence alignments (MSAs) using PRANK+F, a phylogeny-aware progressive aligner distinguishing insertions from deletions (Löytynoja and Goldman, 2005). For each reconstructed MSA, branch lengths on the species tree were re-optimized by maximum likelihood with PhyML v3.3.20211231 (Guindon et al, 2010). Codeml from PAML (Yang, 2007) was used in a few cases where PhyML optimization has failed. Finally, a refined PRANK MSA was inferred for each orthologous group using a species tree with re-optimized branch lengths as a guide tree. The WAG amino acid substitution model (Whelan and Goldman, 2001) was used in all analysis steps, including the ancestral sequence reconstruction described below.

**Fig. 13.**
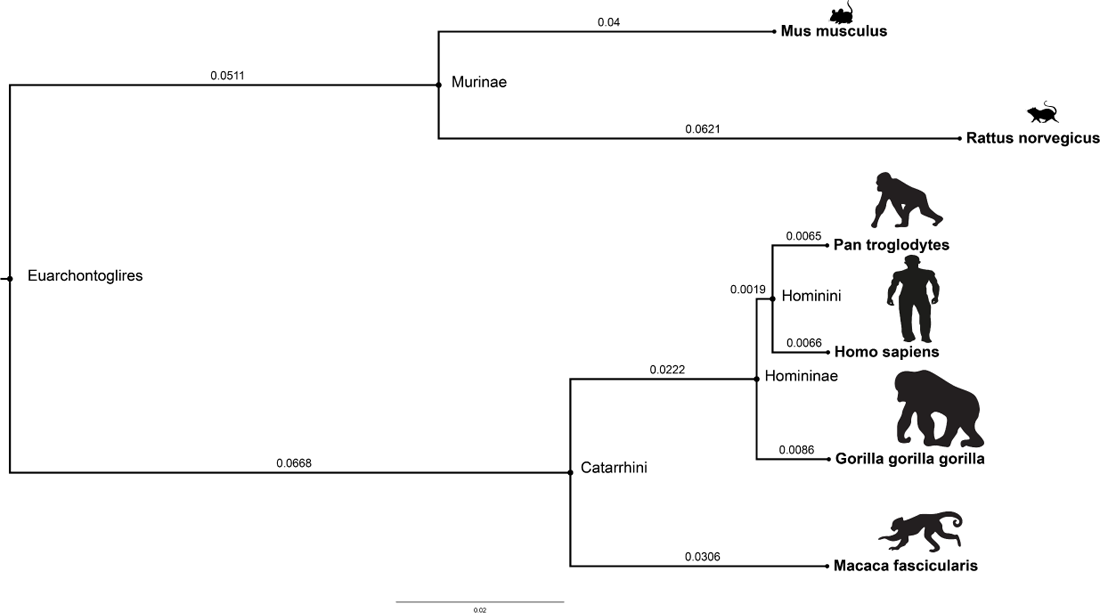
Illustration of the guide tree extracted from 43 *eutherian* mammals. The branch lengths were estimated using pairwise MSA in Ensembl Compara v.105.

### 5.2 Ancestral sequence reconstruction

The refined MSA was used to infer ancestral sequences at all species tree nodes with optimized branch lengths with our recent method implemented in ARPIP (Jowkar et al, 2023). Evolutionary changes on a phylogeny are described via the PIP model (Bouchard-ôté and Jordan, 2013), where insertions follow a Poisson process, while substitutions and deletions follow a continuous-time Markov model with an absorbing state. The ARPIP method includes two main steps. First, the method infers the most probable indel scenario on a given phylogeny, independently for each MSA column. Next, similar to FastML (Pupko et al, 2000), ancestral characters are reconstructed on a subtree of the given phylogeny obtained by pruning it to the inferred indel scenario. For ASR analyses, the root was placed on the internal branch connecting the *Catarrhini* and *Murinae* clades. Then, midpoint rooting was used to define the location of the root on this branch.

### 5.3 Simulating data

We simulated 1000 data sets with INDELible (Fletcher and Yang, 2009). To set realistic parameters, we sampled uniformly at random 1000 OMA groups and extracted the corresponding PRANK MSAs and species trees with PhyML-optimized branch lengths (as described above). For each sample, we simulated a replicate on the PhyML tree using a sequence of 1000 amino acids at the root. We use a Zipfian indel length distribution with *α* = 1.7, a maximum indel length equal to the maximum gap length of the OMA group in question, and an indel rate of 0.1. Sequence lengths in the simulated samples ranged between 336 and 1730 amino acids, while the gap lengths ranged from 1 to 1451 characters. Around 1% of simulations produced biologically unrealistic sequences with extremely long gaps, for example, the sample with a 1451 character long gap. Such samples would be considered noisy in real datasets (possibly due to sequencing errors) and were thus also removed from the simulation analysis before evaluating reconstruction performance. Only four simulated samples contained no gaps at all and were also removed from analysis. The final simulated dataset contained 786 to 1371 amino acid long sequences and the gap lengths ranged from 1 to 235 characters. We provided the true MSA from the simulation and the PhyML tree (i.e. true tree) to ARPIP for ancestral reconstruction.

## 7 Available data and scripts to study indel pattern

This manuscript is accompanied by the scripts used to produce the results. The experimental data used in this manuscript is freely available from https://doi.org/10.5281/zenodo.10798097. The Python scripts used for data processing and analysis are also available at https://github.com/acg-team/ single-char-indel-ASR-preserves-long-indels.

## 8 Ethics declarations

### 8.1 Ethics approval and consent to participate

Not applicable.

### 8.2 Consent for publication

Not Applicable.

## Appendix A Tables and figures

**Table A1.** ARPIP performance in simulation. All metrics include the root sequences and have been computed for each sample individually. We report the averages over the samples.

| Metric | Consistency (%) |
| --- | --- |
| Proportion of correctly inferred ancestral characters | $97.88 \pm 2.01$ |
| Proportion of correctly inferred ancestral columns | $90.35 \pm 2.01$ |
| Proportion of correctly inferred ancestral amino acids (i.e., excluding gaps) | $97.75 \pm 2.55$ |
| Gap precision | $94.27 \pm 5.37$ |
| Gap recall (sensitivity) | $96.99 \pm 3.97$ |
| Gap F-score | $95.46 \pm 3.37$ |
| Gap specificity | $99.29 \pm 1.17$ |

**Table A2.** ARPIP performance in gap character inference by simulation. Performance is shown individually for each internal node.

| Lineage/Clade | Gap consistency/accuracy (%) |  |  |
| --- | --- | --- | --- |
|  | Precision | Recall | F-score |
| Murinae | $98.74 \pm 5.24$ | $99.93 \pm 1.10$ | $99.31 \pm 1.60$ |
| Hominini | $99.99 \pm 0.11$ | $99.9995 \pm 0.02(100\%)$ | $99.9970 \pm 0.06(100\%)$ |
| Homininae | $99.98 \pm 0.20$ | $99.94 \pm 1.51$ | $99.95 \pm 0.97$ |
| Catarrhini | $98.48 \pm 1.75$ | $99.96 \pm 0.60$ | $99.71 \pm 0.99$ |
| Euarchontoglires | $77.71 \pm 17.22$ | $85.49 \pm 17.46$ | $79.44 \pm 15.07$ |

### A.1 Tables related to the accuracy of reconstruction on the simulated data

We report the average accuracy over all the samples.

**Fig. A1.**
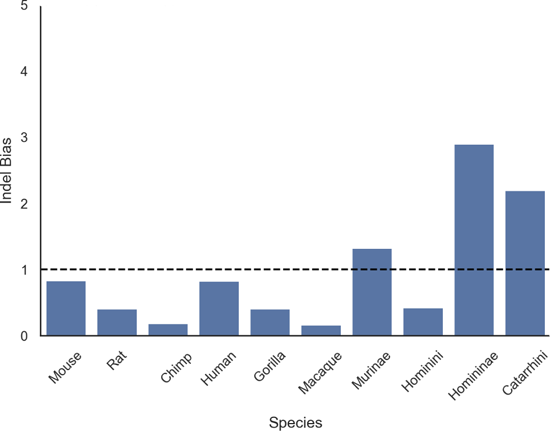
Indel bias (ratio of insertion to deletion events) in mammalian data. A ratio of less than one indicates a bias toward deletions.

### A.2 Indel bias plots for the mammalian and simulated data

**Fig. A2.**
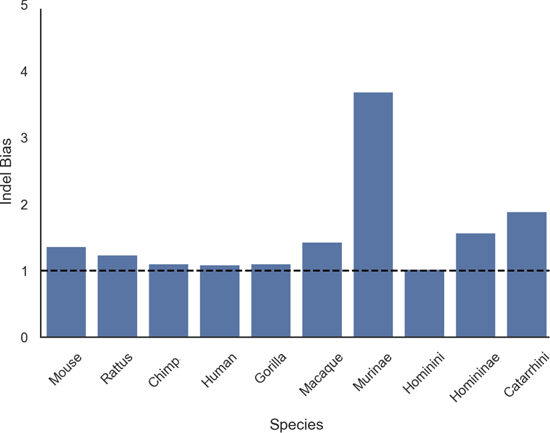
Indel bias (ratio of insertion to deletion events) in simulated data. A ratio of less than one indicates a bias toward deletions.

## Appendix B Study of example reconstructions on simulated data

To get a better intuition for the performance of indel reconstruction under PIP, we have selected two samples from the pool of simulated data for closer examination. The sample *s*_1_ is among the data with the lowest gap retrieval performance, while the second sample *s*_2_ is a sample with a relatively good gap retrieval score.

### B.1 Sample 1: Sub-optimal performance

We have selected two samples from the pool of simulated data to study the performance of gap reconstruction of ARPIP. Sample *s*_1_ is among the samples with the lowest F-score. For *s*_1_, the F-score is 72.86%, while precision and recall are 100% and 57.31%, respectively. This means that all the inferred gaps were correct, but only around half of the gap characters were inferred. The inference accuracy at the root was the lowest not only in this sample but also in all the samples from the simulated dataset (see Tab. A2). Fig. B4 visualizes a segment of *s*_1_ to investigate ASR performance and gap patterns.

**Fig. B3.**
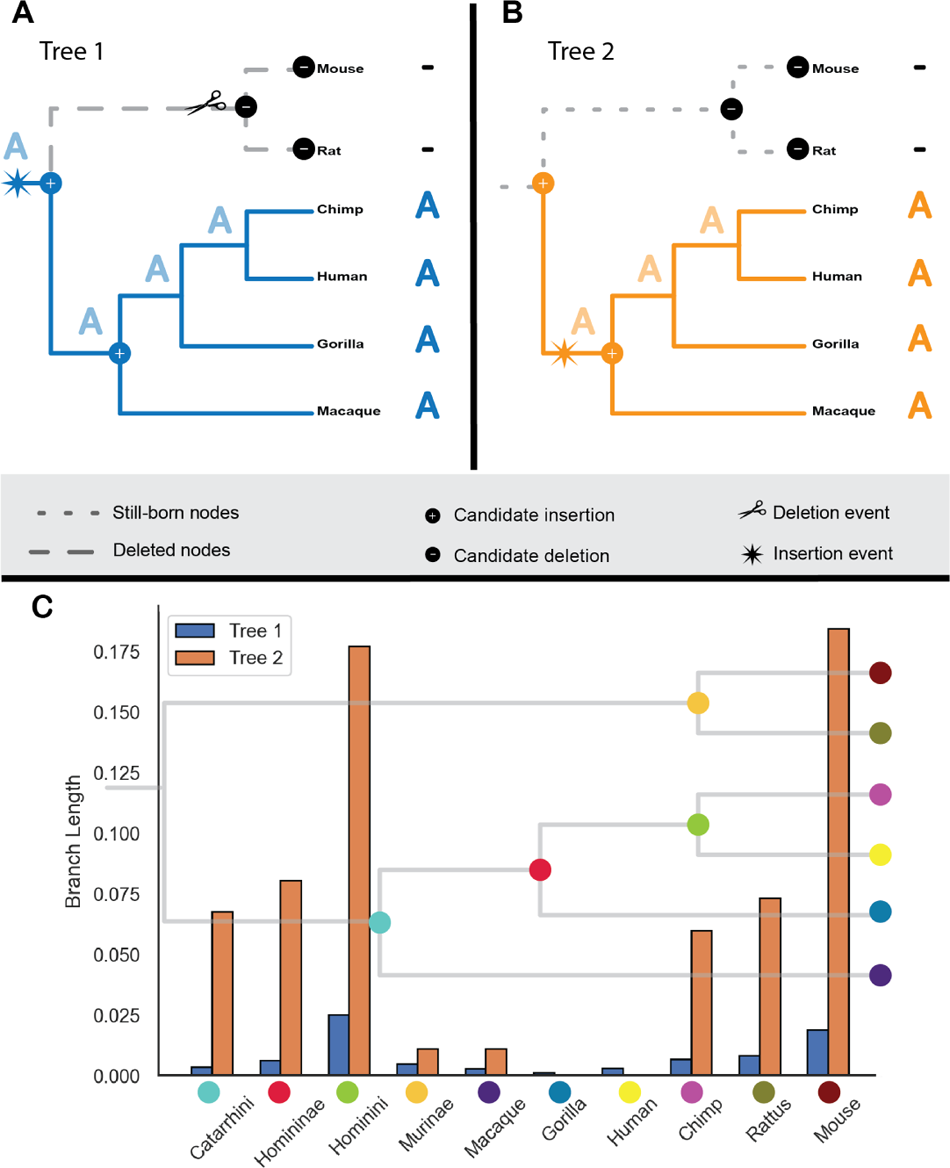
A, B) Two different indel scenarios for a single MSA with various branch lengths. C) Histogram of branch lengths of two selected simulated samples.

**Fig. B4.**
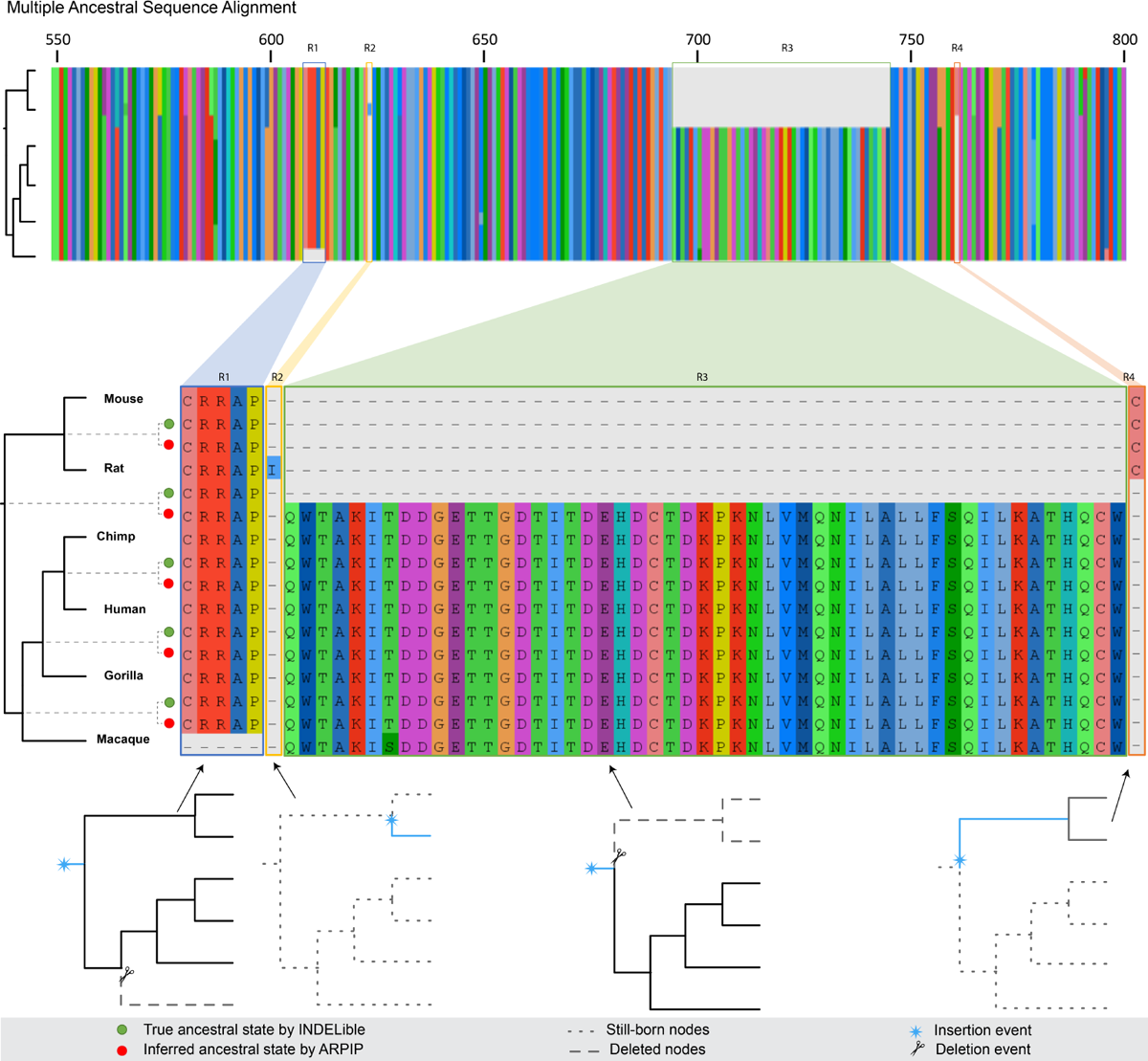
Multiple ancestral sequence alignment of ARPIP inference and INDELible true ancestral states for sites 550 *−* 800 of sample *s*_1_. The indel inference for each site is shown at the bottom of the figure.

**Fig. B5.**
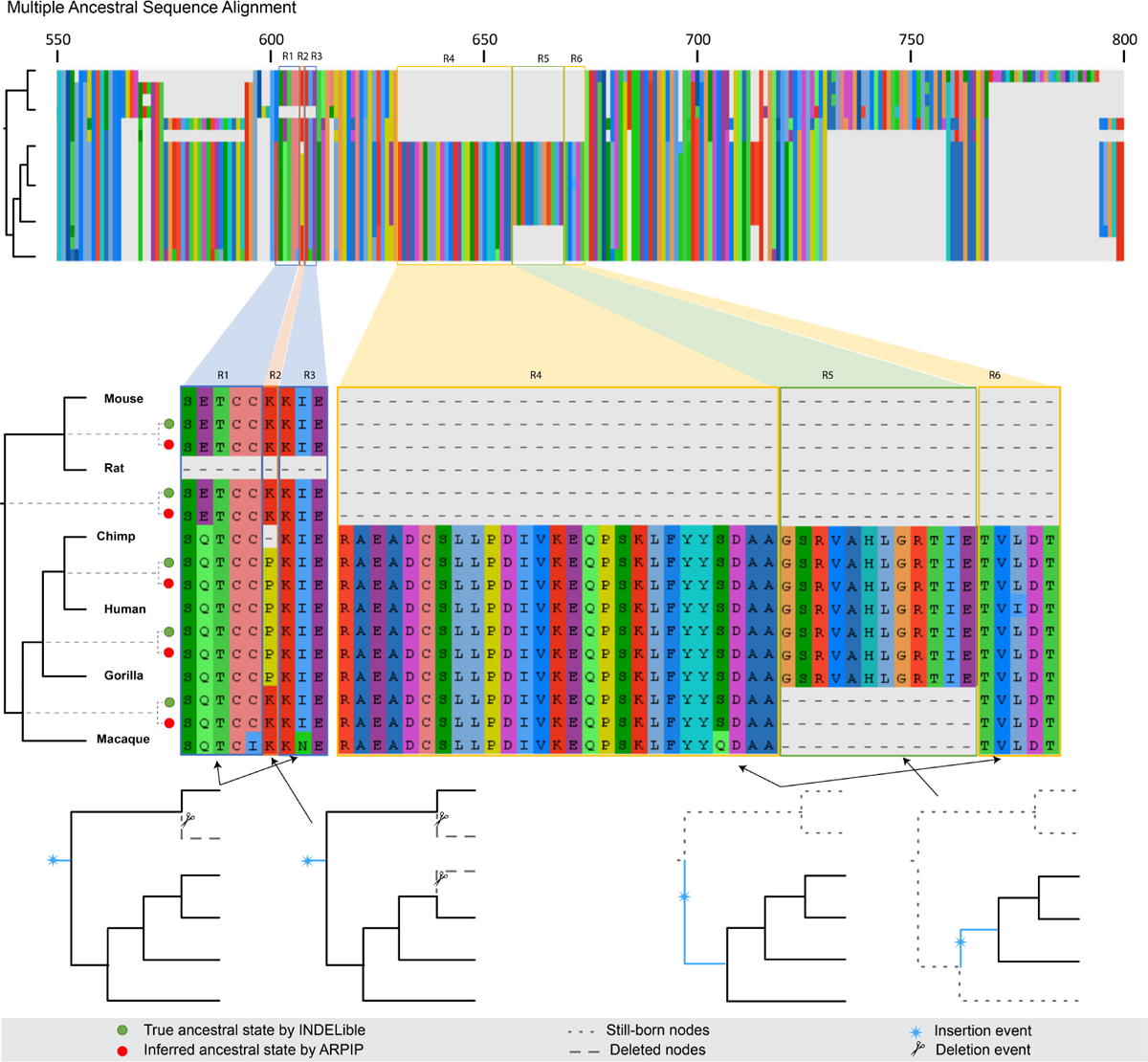
Multiple ancestral sequence alignment of ARPIP inference and INDELible true ancestral states for sites 550 *−* 800 of sample *s*_2_. The indel inference for each site is shown at the bottom of the figure.

Fig. B4 highlights the inferred and true ancestral sequences for four regions of interest. Region R1 depicts five independent insertion and deletion events. Each insertion happened at the root, followed by deletion at the *macaque* taxon. Region R1 does not affect the ancestral gap length distribution, but this typical case happens for a single stretch of gaps at the taxa node. Similarly, region R2 occurs when a single residue in an MSA column exists. A single insertion at the taxa node usually represents a single residue insertion event. This inserted site will show up as a gap in all ancestral nodes, affecting the gap length distribution at the ancestral node, while in reality this site never existed in the ancestor.

Region R3 contains multiple long gaps within both ancestral and taxa species. In this case, as the neighboring site across both ancestral and descendent nodes has the same gap pattern, ARPIP infers the same indel scenario given fixed model parameters. A single insertion at the tree’s root is followed by deletion at the *Murinae* branch.

In *s*_1_, the gap reconstruction accuracy in the root node is very low due to low recall, meaning that ARPIP reconstructs a small fraction of the gaps in the root node. The cause for low gap character retrieval rates remains to be explained. Fig. B3 shows different indel scenarios for a constant MSA column with respect to the branch length of the tree.

Region R4 is a masking indel event of region R3, as we have an insertion event at the branch leading to the node *Murinae*. This is a single-site indel event affecting the ancestral and descendant gap distribution. Notice that we have a single insertion at node *Murinae* without any deletion events.

### B.2 Sample 2: Optimal performance

In addition, we have selected sample *s*_2_ with an overall F-score of 91.36%, resulting from 84.10% precision and 100% recall. This implies that all the gaps were inferred correctly, while a fraction of non-gap characters were falsely inferred as a gap. Fig. B5 illustrates that ARPIP performs well in inferring ancestral sequences despite the complex gap pattern, with 93.42% overall reconstruction accuracy. Moreover, sample *s*_2_ performs relatively well at the root node compared to sample *s*_1_. Fig. B5 shows the gap pattern in two selected neighboring regions (R1-3) and (R4-6).

The PIP model tends to place the insertion events at the root because the Poisson process initiates at the tree’s root. Regions R1 and R3 have a repeated insertion at the root followed by a single deletion event at the *rat* taxon. A neighboring region denoted by R2 has an additional gap between the regions mentioned. ARPIP can adapt the indel event for this specific site while preserving the gap distribution for the other two regions. The gap pattern in these three regions did not affect the gap distribution of ancestral nodes.

The transition from gap pattern R1 to R2 (sites 606 *−* 607) and R2 to R3 (sites 607 *−* 608) suggests that introducing a new gap in another node would have minimal impact on gap inference. The transition from R4 to R5 (sites 656 *−* 657) or R5 to R6 (sites 668 *−* 669) shows the ARPIP can preserve gap distribution for the long ancestral gaps. The results suggest that ARPIP performs exceptionally on neighboring sites with long gaps but suboptimally at the root.

Neighboring segments R4-R6 show two different indel event patterns. We infer that the R4 and R6 segments have an insertion at the *Catarrhini* node, and the R5 segment has an insertion at *Homininae*, without any deletion events at these sites. These three neighboring regions would affect both the ancestral and descendant gap patterns. Like in sample *s*_1_, region R5 separates R4 and R6 without negatively affecting the gap inference. This example shows that ARPIP is relatively good at preserving gap patterns in the neighboring sites.

## Footnotes

1 The percentage of correctly inferred gap characters among all inferred gap characters

2 The percentage of correctly inferred gap characters among all true gap characters

